## Supplemental Figures and Captions for "Evaluation of the fitness benefit conferred by RNA cis-regulators to *Streptococcus pneumoniae* during infection"

**Supplementary Information:**

**Supplemental Tables:** (Tabs in Excel file)

**Table S1:** Accession numbers for the regulatory RNA and downstream protein used to assess extent of conservation across representative *S. pneumoniae* genomes.

**Table S2:** Composition of the Complete Defined Medium (CDM) used to assess extent of gene expression and growth of mutant strains. Medium is made, filter sterilized and used within 3 days of creation.

**Table S3:** Measured parameters from growth curves in Fig. S3 (doubling time, growth rate, carrying capacity, relative growth rate, and relative carrying capacity) for each individual mutant under +ligand (CDM with no dropouts for all RNAs except 1848_guanine, which was guanine supplemented as noted) and -ligand (CDM lacking specific ligand or ligand precursor as noted) conditions. Statistical significance of difference between each mutant to the $WT^C$ is noted. Growth parameters derived from curves in Fig. 6D, 7D, and 7E.

**Table S4:** Average p-values and proportion of samples with p-values <0.05 from data aggregation and sampling analysis.

**Table S5:** Primers used for strain construction and qPCR.

**Supplemental Figures:**

**Figure S1:** Genes regulated by each RNA cis-regulator chosen for study in *S. pneumoniae* TIGR4 categorized by the type of regulator. Locus tags and gene names provided where available. Gene function (transport, biosynthesis, regulatory, unknown) indicated by the color of the box. Operon structure determined in a previous sequencing study [64]. #This regulator was assessed as part of a previously published study [64].

**Figure S2: Regulatory RNA putative secondary structures and mutant gene expression.** (**A-J**): Putative secondary structures are derived from aptamer consensus folding [65] and minimum free energy calculations [95]. Mutations are indicated on each structure. Some mutations are large deletions where the deleted bases are bracketed by an appropriate color bracket (Ligand Insensitive = yellow, ON = green, OFF= red). β-galactosidase activity measured via Miller assay [92], and error bars represent standard deviation and individual biological replicates are indicated by points. Significance of mutant changes in activity upon ligand binding determined via one-way ANOVA followed by Sidak's multiple comparisons test to compare values in the + and – ligand conditions for each mutant. (*$p<0.05$, **$p<0.01$, ***$p<.001$). WT samples are duplicated from Fig. 1A for reference. (**A-D**) TPP riboswitch structures and mutants' β -galactosidase activity in CDM lacking thiamin and including thiamin (-/+ 0.3 μg/ml thiamin). (**E, F)** FMN riboswitch structures and mutants' β -galactosidase activity in CDM (-/+140 ng/mL riboflavin). (**G**) Glycine riboswitch structure and mutants' β -galactosidase activity in CDM (-/+75 μg/mL glycine). % indicates values that are significantly different from one another, but not considered ligand responsive due to the direction of the response. (**H**) Tryptophan T-box structure and mutants' β -galactosidase activity in CDM (-/+204 μg/mL tryptophan). (**I**) Guanine riboswitch structure and mutants' β -galactosidase activity in CDM and CDM supplemented with 50 ug/mL guanine. (**J**) PyrR element preceding SP_0701 structure and mutants' beta-galactosidase activity in CDM (-/+ 20 μg/mL uracil) (**K)** PyrR element preceding SP_1286 and qPCR measurement of gene expression. Individual points indicate biological replicates (average of 2 technical replicates).

**Figure S3: Growth Curves for regulatory RNA mutants.** Growth curves in a complete synthetic medium (CDM) in the presence and absence of ligand. All curves represent at least two biological replicates. Error bars represent standard error across all replicates (n = 5-9). Doubling time and carrying capacity measurements extracted from each individual curve under each condition are indicated on graphs below. In each condition mutants were compared to the $WT^C$ via Kruskal-Wallis test followed by Dunn's test of multiple comparisons, those displaying significant changes are indicated ((*p<0.05, **p<0.01, ***p<0.001, ****p<0.0001) **(A)** 0716_TPP mutants grown in the presence and absence of thiamin. **(B)** 0719_TPP mutants grown in the presence and absence of thiamin. **(C)** 0726_TPP mutants grown in the presence and absence of thiamin. **(D)** 2199_TPP mutants grown in the presence and absence of thiamin. **(E)** 0178_FMN mutants grown in the presence and absence of riboflavin #. **(F)** 0488_FMN mutants grown in the presence and absence of riboflavin#. **(G)** 0701_pyrR regulator mutants grown in the presence and absence of uracil#. **(H)** 1286_pyrR mutants grown in the presence and absence of uracil#. **(I)** Growth curves for 0408_Glycine mutants grown in the presence and absence glycine. **(J)** Growth curves for 1069_TrpT-box mutants grown in the presence and absence tryptophan. **(K)** 1847_Guanine mutants grown in the presence and absence of guanine. (**L**) Parameters extracted from previously published growth curves for the 1272_pyrR mutants [64] **(M)** Growth curves in a semi-defined minimal media (SDMM) for 0178_FMN_WT$^C$ and mutants showing autolysis death phase of characteristic of *S. pneumoniae* growth in richer medium. #These growth curves are also shown in Fig. 6 or Fig. 7, but repeated here for accessibility to the entire data set.

**Figure S4: Summarized Relative fitness and relative growth parameters for individual mutant strains.** The fitness mean of each individual strain is displayed for nasopharynx colonization **(A)**, lung infection **(B)**, and transition to blood models **(C)**. Orange circles = mLI (Ligand Insensitive), green triangle=mON (constitutively active gene expression), and red square =mOFF (repressed gene expression), error bars correspond to the standard deviation. Filled points correspond to strains with a statistically significant change (p. adj <0.05) from the *in vitro* competition (rich medium) control (Kruskal-Wallis test followed by Dunn's test of multiple comparisons, adj p<0.05). Open points are not significantly distinct from the *in vitro* control competition. The black-filled point (1286_pyrR_mON) corresponds to an environment under which none of the mutant strain was recovered despite repeated attempts, indicating a fitness close to 0. Graphs representing individual mouse competitions for each strain are found in Fig. S5. Relative carrying capacity **(D,E)** and relative growth rate **(F,G)** for mutant strains compared to the WT$^C$ strain. Points represent the mean of 5-9 replicates, and the error bars represent the standard deviation. Some error bars are smaller than the size of the point and therefore not visible. Open points are not statistically significantly different from the WT$^C$ strain. Colored points represent values that are statistically significant from the WT$^C$ strain (Kruskal-Wallis test followed by Dunn's multiple comparisons test, adj. p <0.05, Fig. S4, Table S3). Vertical lines separate groups of riboswitches interacting with the same ligand.

**Fig. S5: Fitness values for individual mouse lung infections and nasopharynx colonization assays compared to the *in vitro* control in semi-defined or rich medium.** Each data point consists of a fitness value determined from a single mouse infection, or *in vitro* competition in rich medium. Statistical difference from the control experiment determined using the Kruskal-Wallis test followed by Dunn's test of multiple comparisons. (*=adj. p<0.05, ** adj. p<0.01, *** p<.001, **** adj. p<.0001). **(A)** 0719_TPP WT$^C$, mLI, mON and mOFF. **(B)** #0178_FMN WT$^C$, mLI, mON and mOFF. **(C)** #0488_FMN WT$^C$, mLI, mON and mOFF. **(D)** #0701_pyrR WT$^C$, mLI, mON and mOFF. **(E)** #1286_pyrR WT$^C$, mLI, and mON. **(F)** 0408_Glycine WT$^C$, mLI, mON and mOFF. **(G)** 1069_Trp-Tbox WT$^C$, mLI, mON and mOFF. **(H)** 1847_Guanine WT$^C$, mLI, mON and mOFF. #These graphs are also shown in Fig. 6 or Fig. 7, but repeated here for accessibility to the entire data set

**Fig. S6: Modelling co-culture indicates infection fitness defects cannot be directly attributed planktonic growth parameters. (A)** Fitness modelled using a variety of parameter sets as indicated to demonstrate sensitivity of the modelled fitness to parameters. **(B)** Modelled fitness during exponential phase (120 minutes) and **(C)** stationary phase (500 minutes) for all RNA regulator mutants based on their relative growth rate and carrying capacity in CDM lacking the target nutrient as reported on Table S3. Modelled fitness calculated based on a co-culture model with the growth rate of the reference strain $r_a$=0.0252 and carrying capacity $K_a$=0.462. Growth rate and carrying capacity of test strains are scaled by the relative growth rate and carrying capacity such that $r_b$=$r_a$*relative growth rate, and $K_b$= $K_a$*relative carrying capacity on Table S3. Horizontal line drawn at fitness $W$ = 1 for visual reference. Error was estimated by repeating calculations with values of $r_a$ and $K_a$ adjusted by the standard deviations of the mean reported on Table S3. **(D)** Negative control *in vitro* competitions conducted between *S. pneumoniae* TIGR4 and 0701_pyrR_WT$^C$ or 1278_pyrR_WT$^C$ show no significant change in fitness. Bars represent mean fitness and error bars standard deviation for individual biological replicates shown as points.

**Fig. S7: Modelled fitness correlates weakly with measured fitness from *in vivo* environments compared to correlations between *in vivo* fitness measures. (A-C)** Modelled stationary phase fitness vs. mean measured fitness in the nasopharynx, lung, and blood. **(D-F)** Modelled exponential phase fitness vs. mean measured fitness in the nasopharynx, lung, and blood. **(G-H)** Comparison of the three different *in vivo* environments shows significant correlation in fitness values. Pearson's correlation coefficient and Bonferroni adjusted p-value (N=9) are reported for each comparison. Error bars correspond to error bars reported in Fig. S4A-C, and S6B, C.

Figure S1

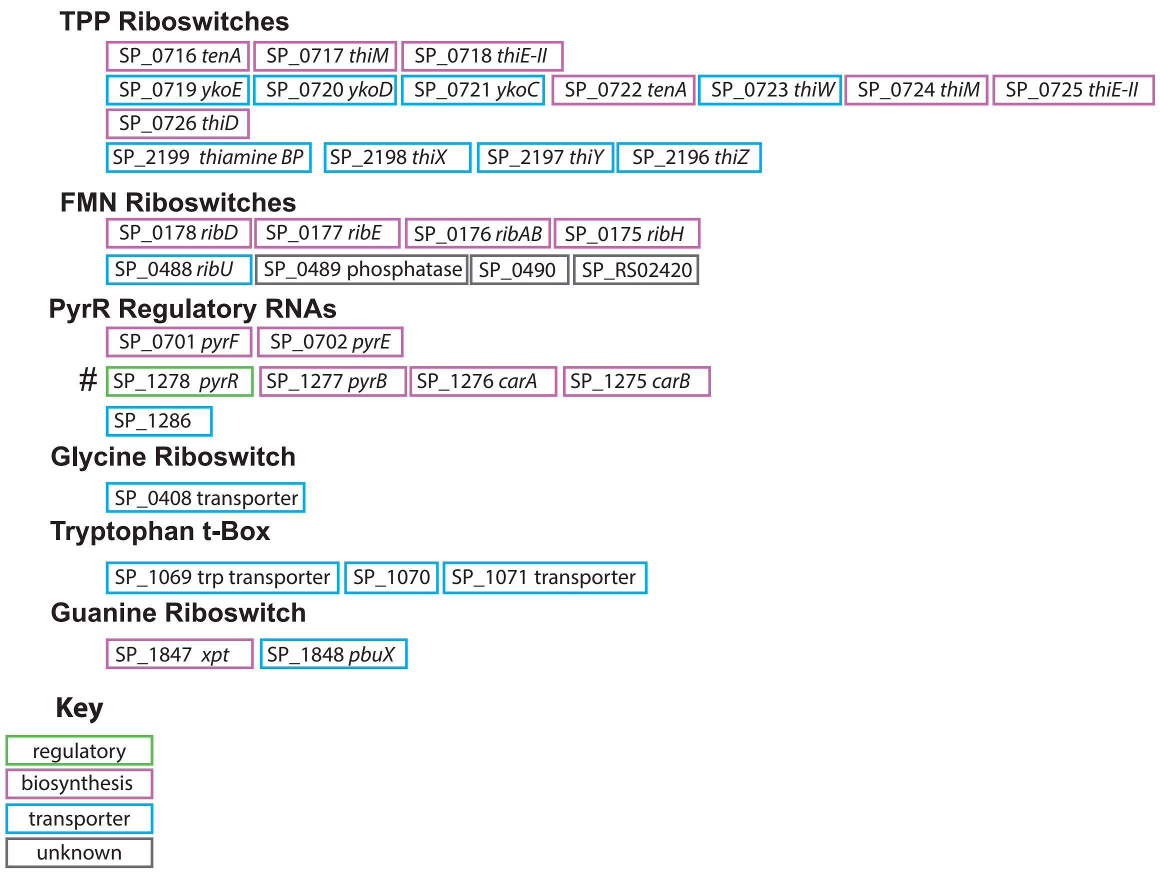
TPP Riboswitches
SP_0716 tenA
SP_0717 thiM
SP_0718 thiE-II
SP_0719 ykoE
SP_0720 ykoD
SP_0721 ykoC
SP_0722 tenA
SP_0723 thiW
SP_0724 thiM
SP_0725 thiE-II
SP_0726 thiD
SP_2199 thiamine BP
SP_2198 thiX
SP_2197 thiY
SP_2196 thiZ
FMN Riboswitches
SP_0178 ribD
SP_0177 ribE
SP_0176 ribAB
SP_0175 ribH
SP_0488 ribU
SP_0489 phosphatase
SP_0490
SP_RS02420
PyrR Regulatory RNAs
SP_0701 pyrF
SP_0702 pyrE
#
SP_1278 pyrR
SP_1277 pyrB
SP_1276 carA
SP_1275 carB
SP_1286
Glycine Riboswitch
SP_0408 transporter
Tryptophan t-Box
SP_1069 trp transporter
SP_1070
SP_1071 transporter
Guanine Riboswitch
SP_1847 xpt
SP_1848 pbuX
Key
regulatory
biosynthesis
transporter
unknown

**Figure S2** pg 1

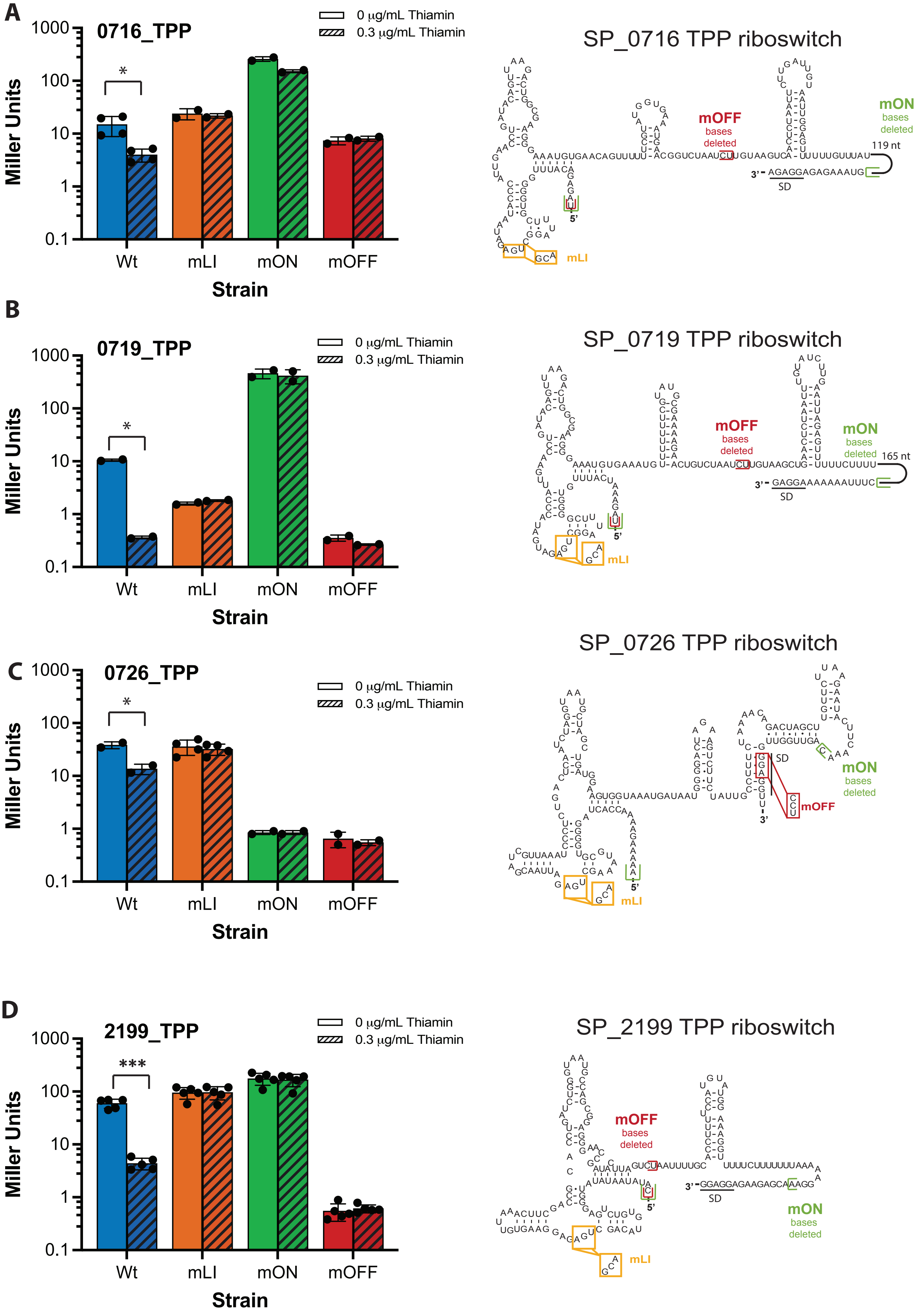

**Figure S2** pg 2

**E**

### SP_0178 FMN riboswitch

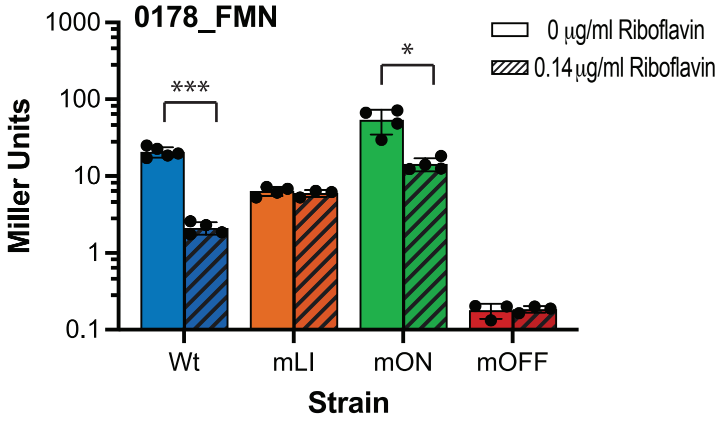


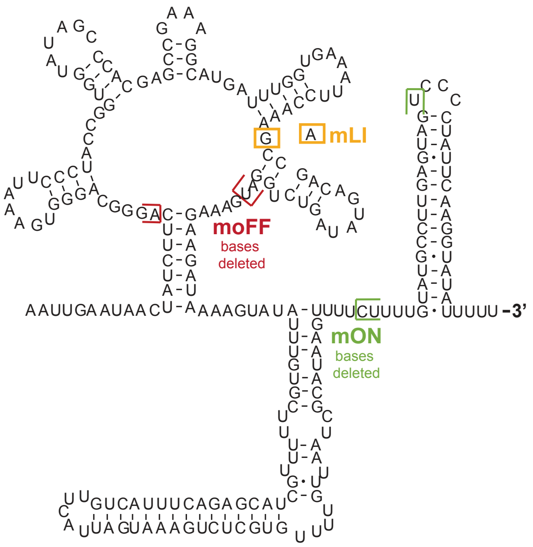


**F**

### SP_488 FMN riboswitch

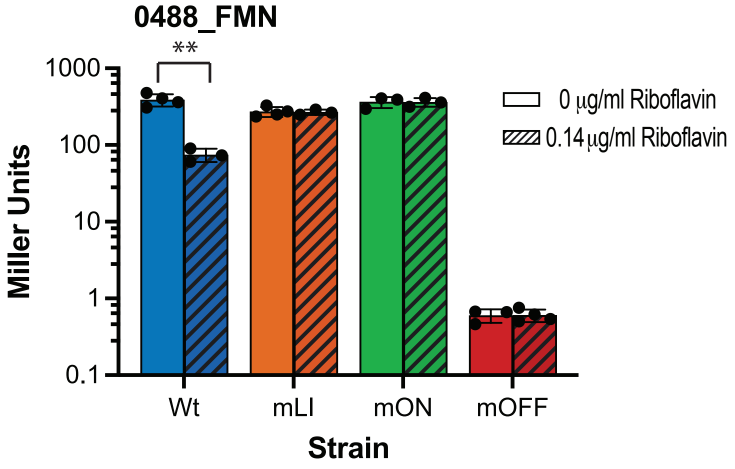


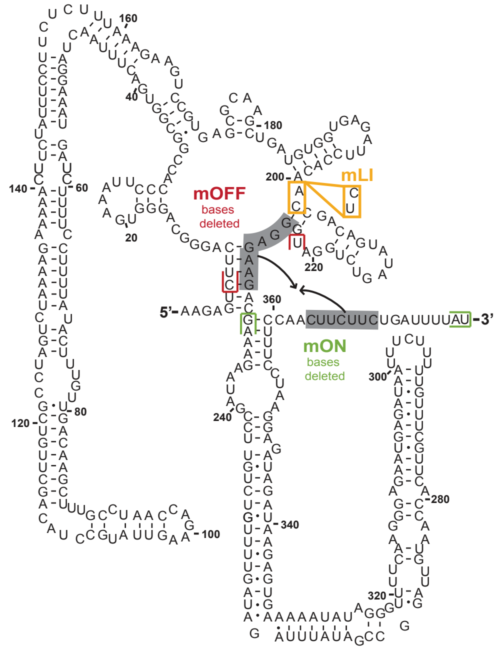

**Figure S2** pg 3

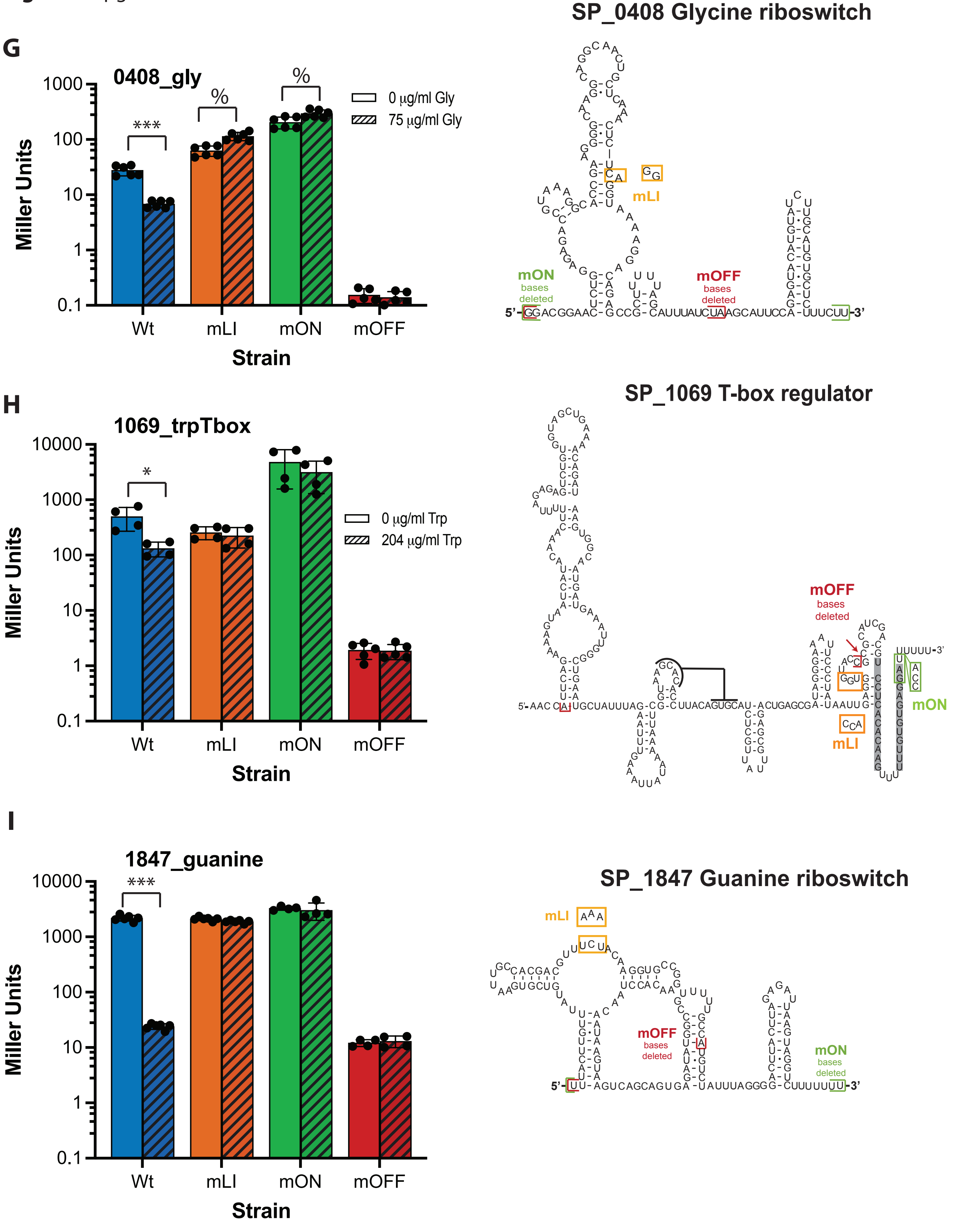

**Figure S2** pg 4

**J**

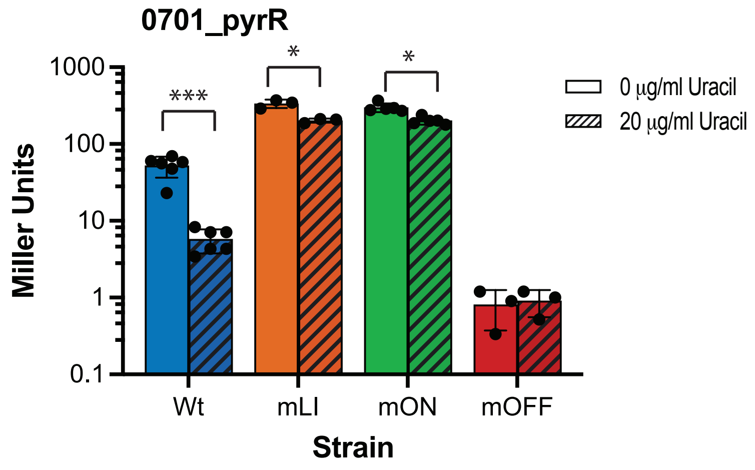


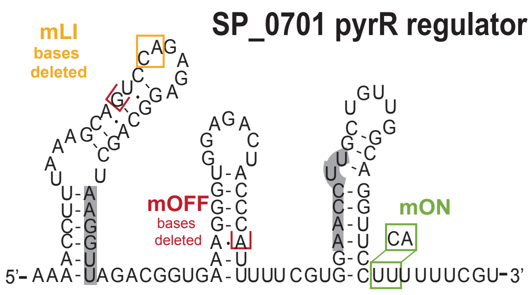


**K**

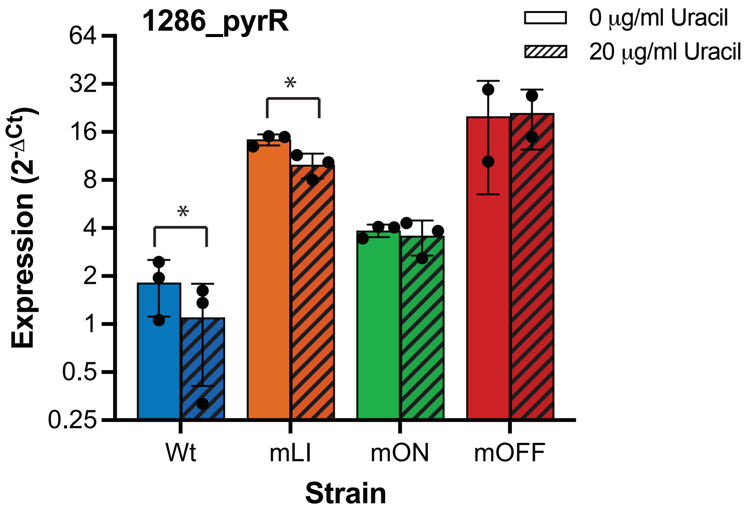


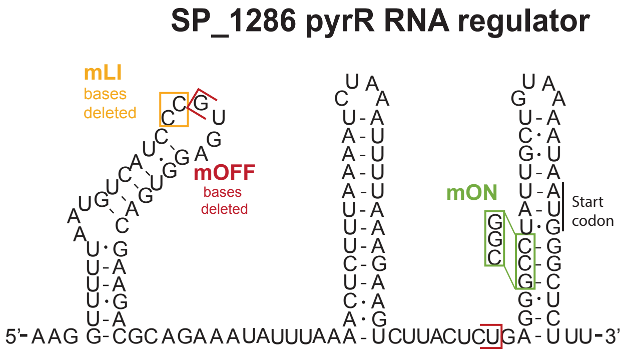

**Figure S3** pg 1

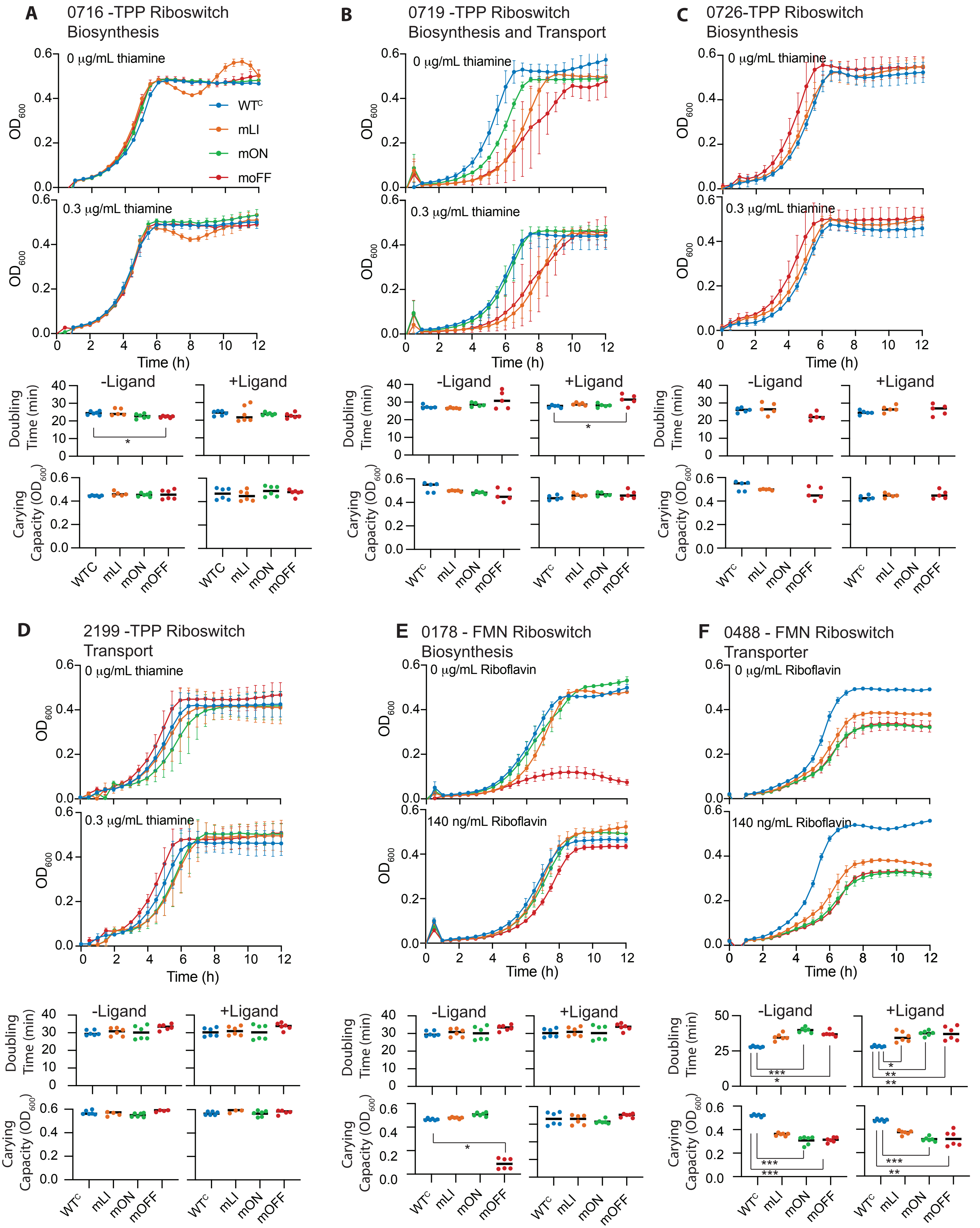

**Figure S3** pg 2

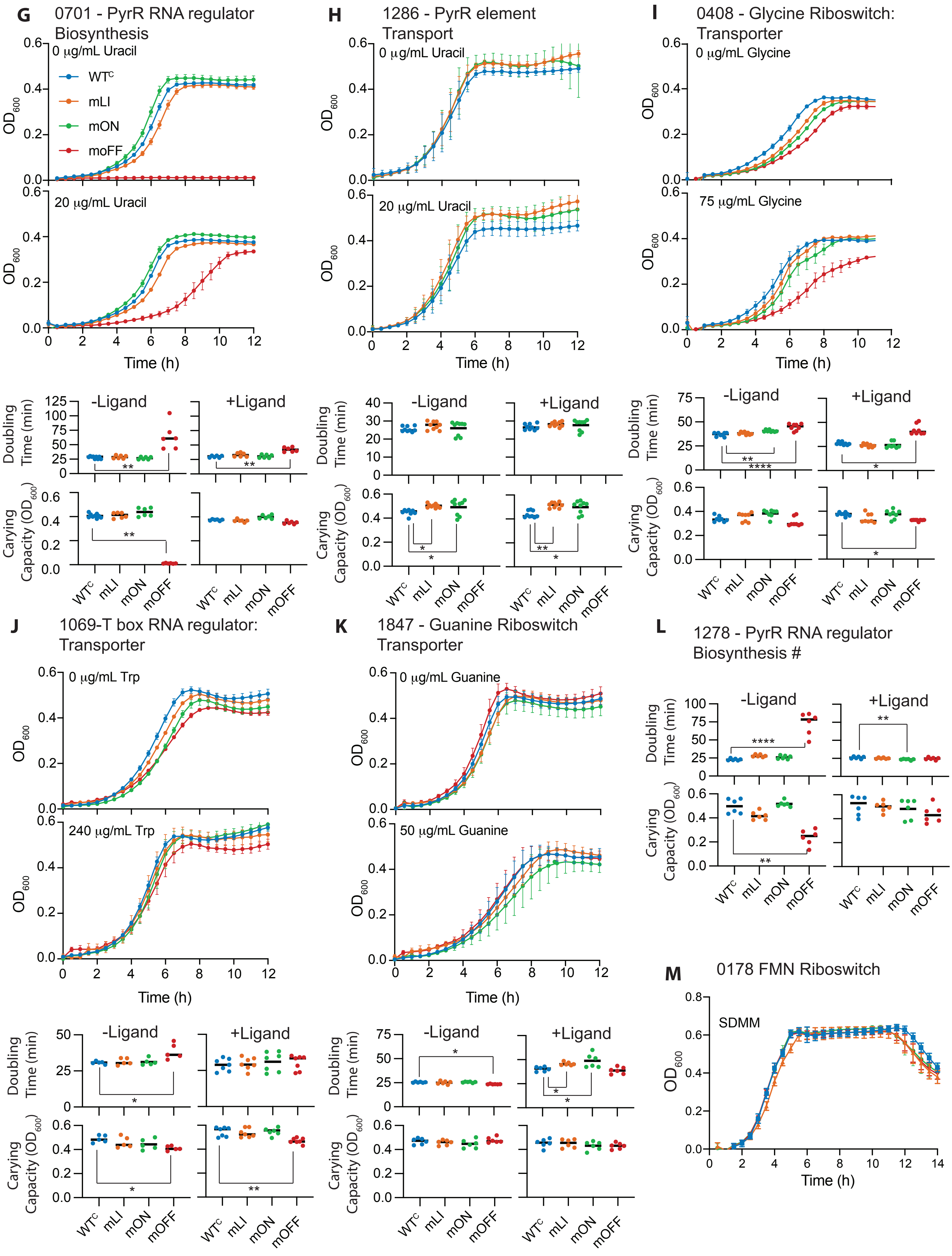

**Figure S4**

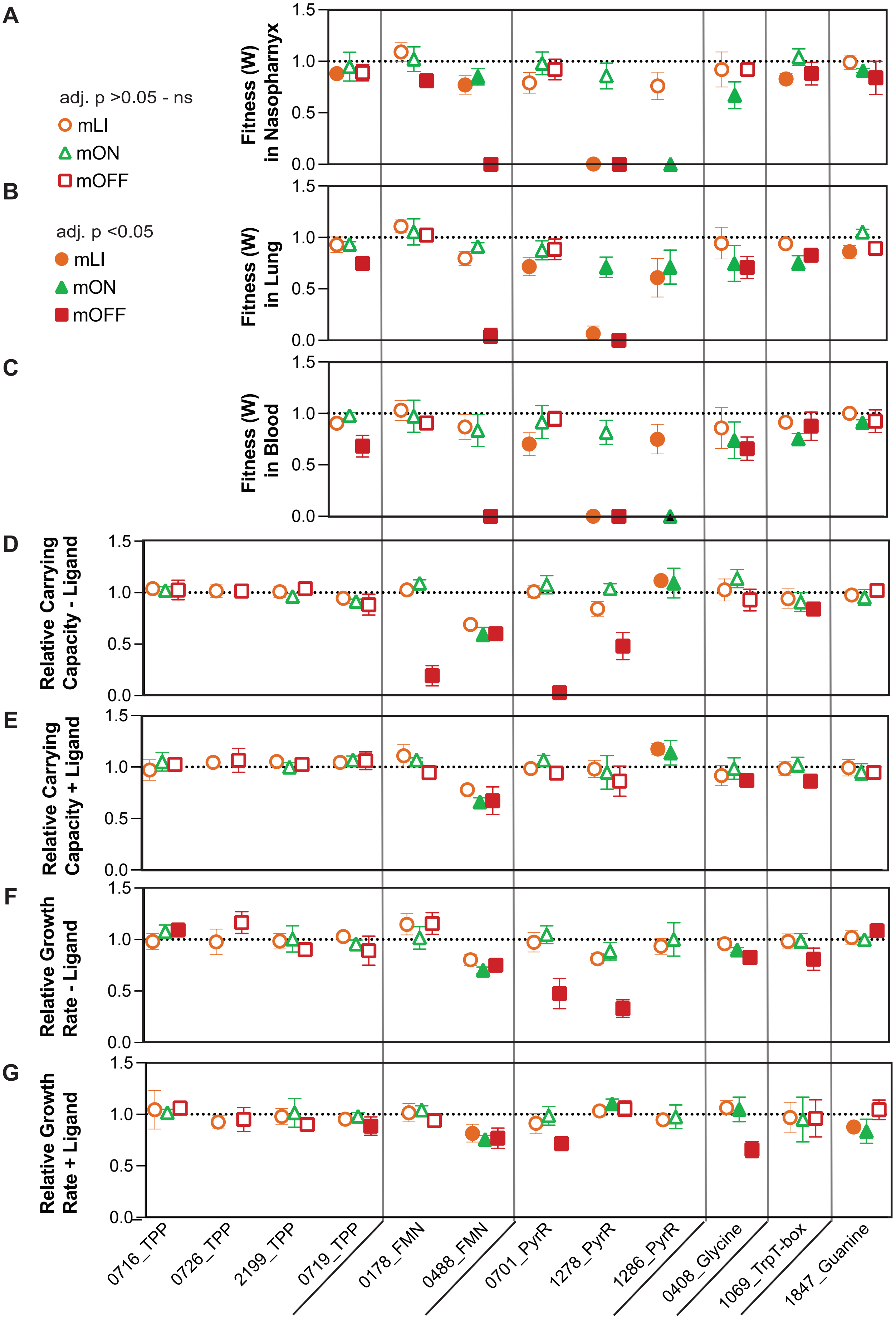
A
B
C
D
E
F
G
adj. p >0.05 - ns
mLI
mON
mOFF
adj. p <0.05
mLI
mON
mOFF
Fitness (W) in Nasopharynx
Fitness (W) in Lung
Fitness (W) in Blood
Relative Carrying Capacity - Ligand
Relative Carrying Capacity + Ligand
Relative Growth Rate - Ligand
Relative Growth Rate + Ligand
1.5
1.0
0.5
0.0
0716_TPP
0726_TPP
2199_TPP
0719_TPP
0178_FMN
0488_FMN
0701_PyrR
1278_PyrR
1286_PyrR
0408_Glycine
1069_TrpT-box
1847_Guanine

**Figure S5** pg 1

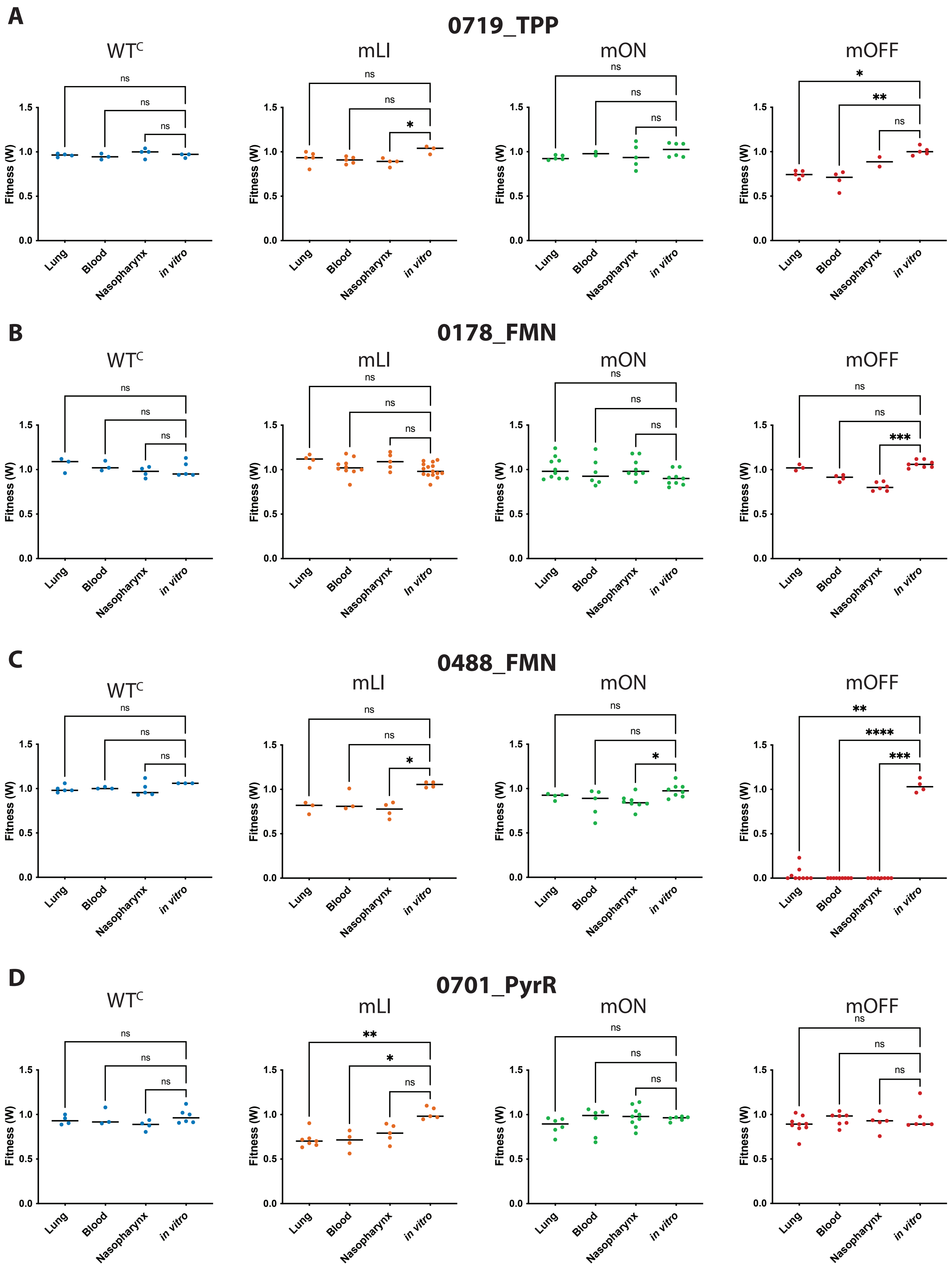

**Figure S5** pg 2

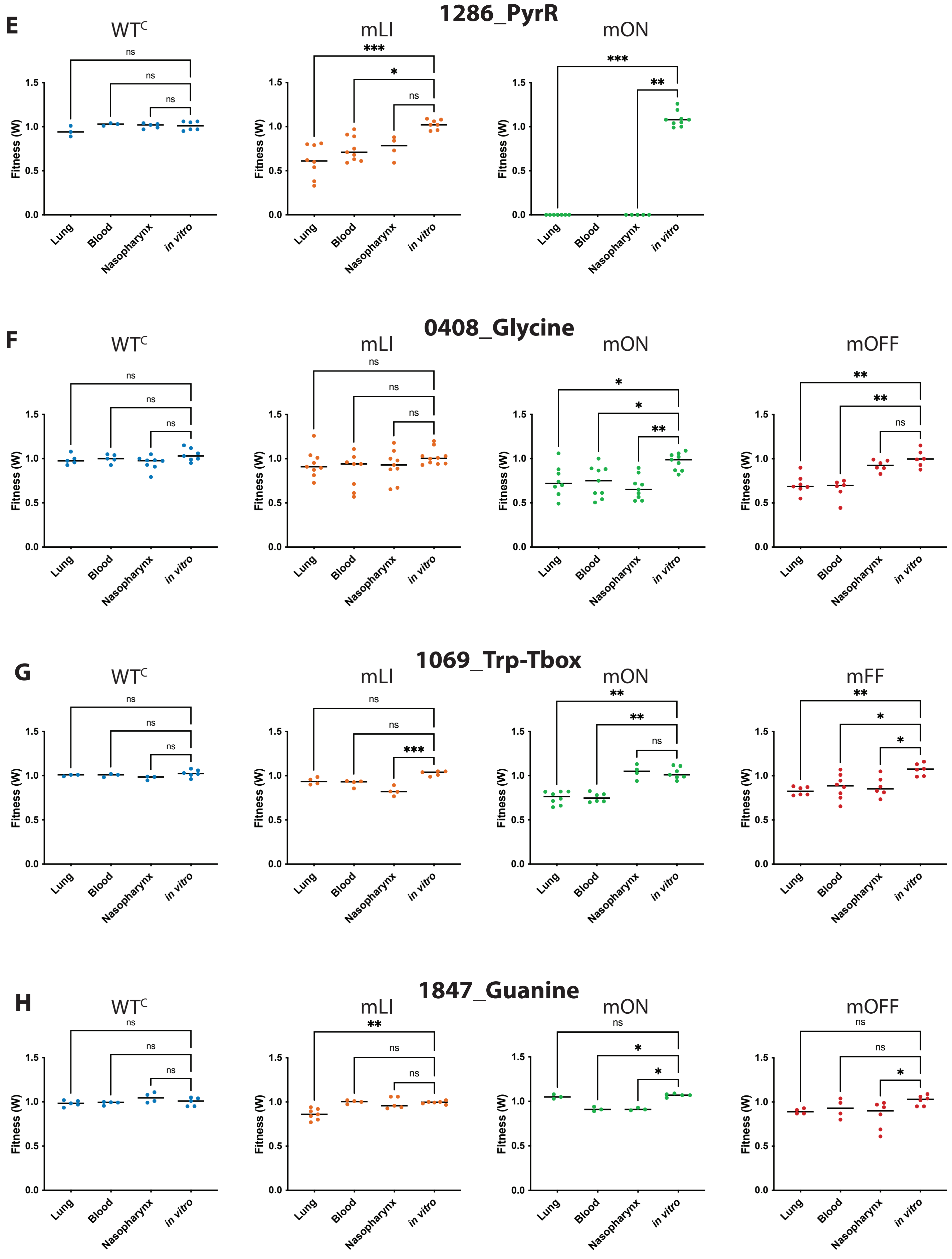

### Figure S6

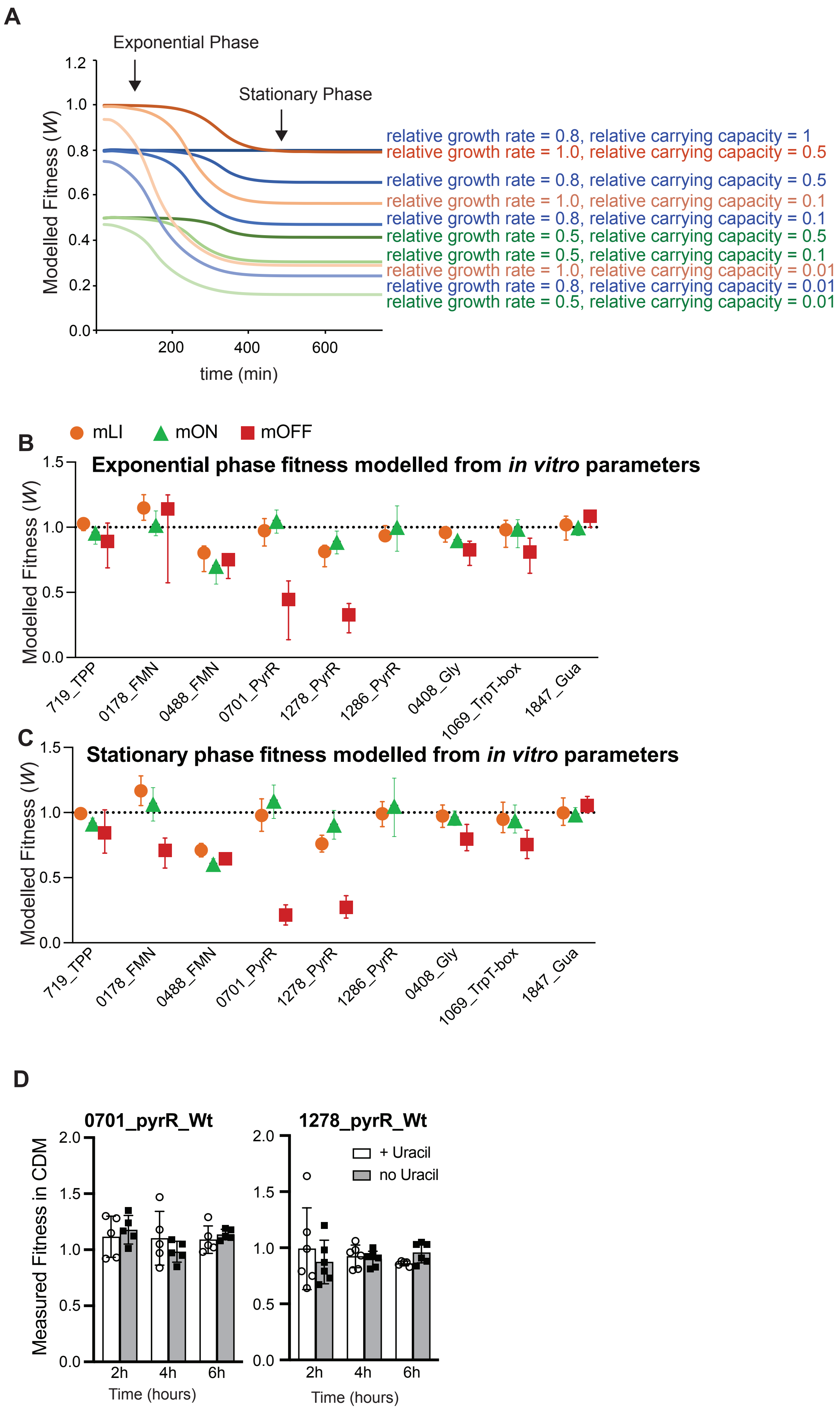
A
Exponential Phase
Stationary Phase
Modelled Fitness (W)
time (min)
relative growth rate = 0.8, relative carrying capacity = 1
relative growth rate = 1.0, relative carrying capacity = 0.5
relative growth rate = 0.8, relative carrying capacity = 0.5
relative growth rate = 1.0, relative carrying capacity = 0.1
relative growth rate = 0.8, relative carrying capacity = 0.1
relative growth rate = 0.5, relative carrying capacity = 0.5
relative growth rate = 0.5, relative carrying capacity = 0.1
relative growth rate = 1.0, relative carrying capacity = 0.01
relative growth rate = 0.8, relative carrying capacity = 0.01
relative growth rate = 0.5, relative carrying capacity = 0.01
B
mLI
mON
mOFF
Exponential phase fitness modelled from in vitro parameters
Modelled Fitness (W)
719_TPP
0178_FMN
0488_FMN
0701_PyrR
1278_PyrR
1286_PyrR
0408_Gly
1069_TrpT-box
1847_Gua
C
Stationary phase fitness modelled from in vitro parameters
Modelled Fitness (W)
D
0701_pyrR_Wt
1278_pyrR_Wt
Measured Fitness in CDM
+ Uracil
no Uracil
Time (hours)

**Figure S7**

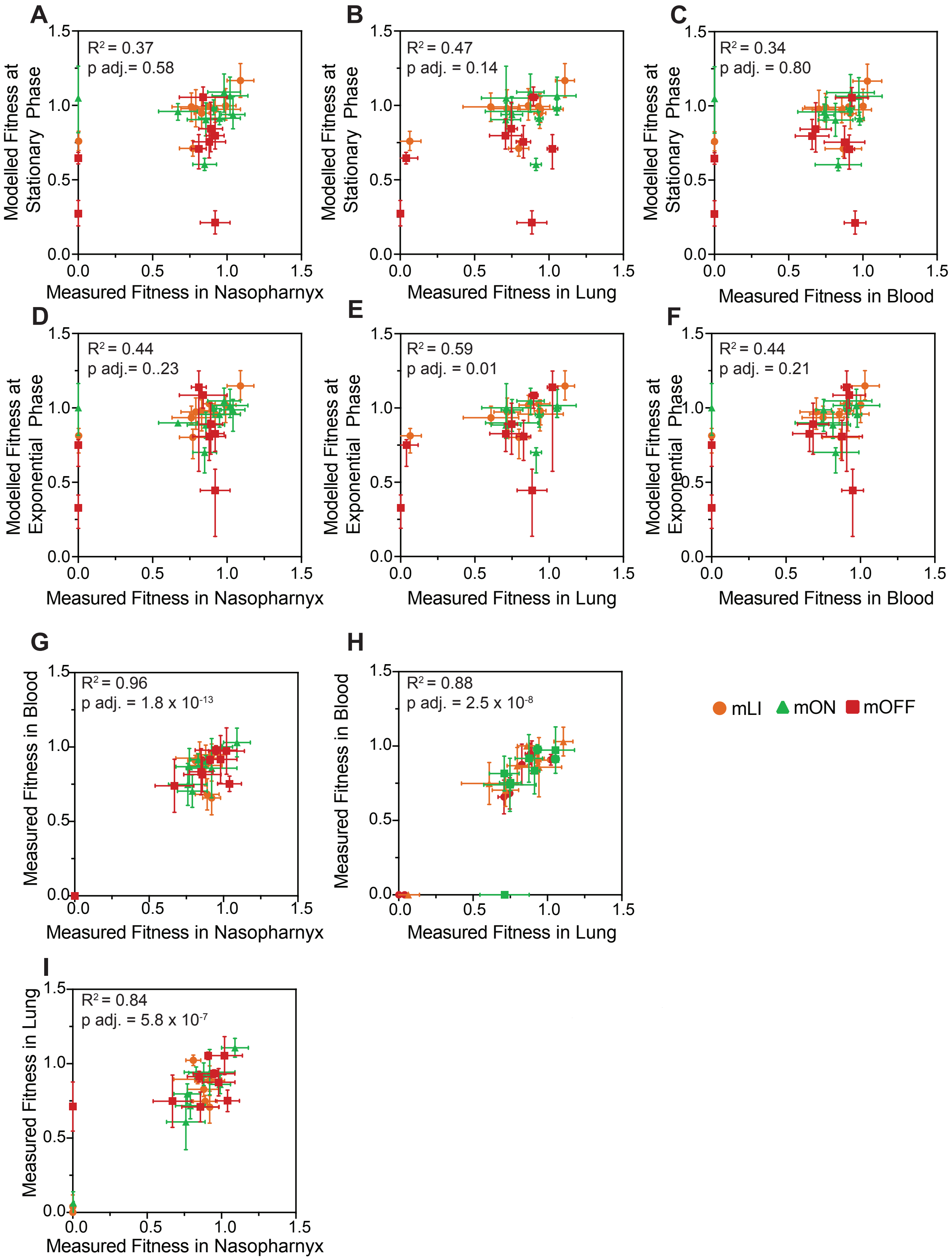

A
R2 = 0.37
p adj.= 0.58
Modelled Fitness at Stationary Phase
Measured Fitness in Nasopharnyx
B
R2 = 0.47
p adj. = 0.14
Modelled Fitness at Stationary Phase
Measured Fitness in Lung
C
R2 = 0.34
p adj. = 0.80
Modelled Fitness at Stationary Phase
Measured Fitness in Blood
D
R2 = 0.44
p adj.= 0..23
Modelled Fitness at Exponential Phase
Measured Fitness in Nasopharnyx
E
R2 = 0.59
p adj. = 0.01
Modelled Fitness at Exponential Phase
Measured Fitness in Lung
F
R2 = 0.44
p adj. = 0.21
Modelled Fitness at Exponential Phase
Measured Fitness in Blood
G
R2 = 0.96
p adj. = 1.8 x 10-13
Measured Fitness in Blood
Measured Fitness in Nasopharnyx
H
R2 = 0.88
p adj. = 2.5 x 10-8
Measured Fitness in Blood
Measured Fitness in Lung
I
R2 = 0.84
p adj. = 5.8 x 10-7
Measured Fitness in Lung
Measured Fitness in Nasopharnyx
mLI
mON
mOFF
